# Molecular mechanism of the Orai channel activation

**DOI:** 10.1101/490516

**Authors:** Xiaofen Liu, Guangyan Wu, Yi Yu, Xiaozhe Chen, Renci Ji, Jing Lu, Xin Li, Xing Zhang, Xue Yang, Yuequan Shen

## Abstract

The Orai channel is characterized by voltage independence, low conductance and high Ca^2+^ selectivity and plays an important role in Ca^2+^ influx through the plasma membrane. How the channel is activated and promotes Ca^2+^ permeation are not well understood. Here, we report the crystal structure and cryo-electron microscopy reconstruction of a *Drosophila melanogaster* Orai mutant (P288L) channel that is constitutively active according to electrophysiology. The open state of the Orai channel showed a hexameric assembly in which six TM1 helices in the center form the ion-conducting pore, and six TM4 helices in the periphery form extended long helices. Orai channel activation requires conformational transduction from TM4 to TM1 and eventually causes the basic section of TM1 to twist outward. The wider pore on the cytosolic side aggregates anions to increase the potential gradient across the membrane and thus facilitate Ca^2+^ permeation. The open-state structure of the Orai channel offers insights into channel assembly, channel activation and Ca^2+^ permeation.

---

Calcium signaling is essential in a broad range of biological processes ^1^. In metazoans, store-operated calcium entry (SOCE) is one of the major extracellular calcium influx pathways not only in excitable cells but also particularly in nonexcitable cells ^2, 3^. The action of extracellular ligands triggers the release of Ca^2+^ from the endoplasmic reticulum (ER). The stromal interaction molecule (STIM) located on the ER senses Ca^2+^ depletion within the ER and, in response, undergoes oligomerization and translocation to the ER-plasma membrane (PM) junction, where it couples with and activates the Ca^2+^-selective channel Orai ^4, 5^. The Orai protein belongs to the family called store-operated calcium channels (SOCs), which have unique features among ion channels, including voltage independence, low conductance and high Ca^2+^ selectivity ^6-8^. In humans, there are three isoforms (Orai1-3) of Orai proteins. They are highly homologous to each other (∼ 62% overall sequence identity) ^5^. Each Orai contains four transmembrane helices with the N-terminal and C-terminal ends located inside the cytosol ^9^. Orai channel-mediated Ca^2+^ signaling plays an important role in multiple physiological processes ^10^. Consequently, loss-of-function mutations and gain-of-function mutations of Orai1 have been identified in human patients and found to cause various diseases, such as severe combined immunodeficiency, skeletal myopathy, Stormorken syndrome, and others ^11, 12^.

The Orai channel is assembled from multiple subunits ^13, 14^. The precise stoichiometry of the Orai channel has been in great debate. Several groups have reported that the Orai channel is a dimer in the resting state and forms a tetramer in the activated state ^15, 16^. The crystal structure of *Drosophila* Orai showed a trimer of the Orai dimer, forming an approximately hexameric assembly ^17^. However, the hexameric stoichiometry of the Orai channel was brought into question by studies of single-molecule photobleaching ^18^ and artificially linked hexameric concatemers of human Orai1 ^19^. Interestingly, recent studies of the concatenated Orai1 channel indicated that the Orai1 channel functions as a hexamer ^20-22^. Therefore, hexameric stoichiometry is generally accepted as one of the major conformations of the Orai channel ^5, 23, 24^.

The current crystal structure of *Drosophila melanogaster* hexameric Orai represents an inactive conformation ^17^. The innermost transmembrane 1 (TM1) from each subunit forms a closed ion pore, and three other transmembrane helices are arranged around TM1. The selectivity filter is presumably formed by a ring of six glutamate residues on the extracellular side of the pore. The mechanism of channel activation and Ca^2+^ permeation remains unclear. Here, we determined the structure of the constitutively active *Drosophila melanogaster* Orai (dOrai) mutant P288L using both X-ray crystallography and cryo-electron microscopy. The open state of the Orai1 structure depicts the mechanism of channel activation and Ca^2+^ permeation.

## Results

### The dOrai-P288L channel is constitutively active

To obtain the three-dimensional structure of constitutively active Orai, a construct of dOrai (hereafter referred to as dOrai-P288L) consisting of amino acids 132-341 with three mutations (C224S, C283T and P288L) was selected and purified (Supplementary Fig. 1). The *Drosophila melanogaster* Orai P288L mutant corresponds to the P245L gain-of-function mutation of human Orai1, which causes overlapping syndromes of tubular myopathy and congenital miosis ^12, 25^. To verify that the dOrai-P288L channel showed constitutive activity, electrophysiology was carried out. First, we transiently transfected HEK-293T cells with a plasmid encoding the GFP-tagged dOrai-P288L channel and showed that the channel was localized to the cell surface (Supplementary Fig. 2a). Next, we used whole-cell patch clamp measurements to record the Ca^2+^ current in HEK-293T cells expressing the dOrai-P288L channel, showing the inward rectifying Ca^2+^ current with a reversal potential close to + 50 mV (**Fig. 1a**). Furthermore, in the absence of divalent cations, we observed clear monovalent cation currents (**Fig. 1b**). The addition of Ca^2+^ significantly inhibited monovalent cation currents (**Figs. 1b, 1c**). Finally, we used single channel current recording *in vitro* to verify the activity of the purified dOrai-P288L channel. The monodisperse and high-purity protein (Supplementary Fig. 2b) was reconstituted into a planar lipid bilayer, as reported by our previous research ^26^. No current signal was observed at −100 mV without the addition of the channel protein. After the addition of the channel protein, obvious inward Ca^2+^ currents of approximately 2 pA at −100 mV were recorded (**Fig. 1d**). The single-channel Ca^2+^ currents gradually diminished as the stimulation potential depolarized from −100 mV to 0 mV. Additionally, the single-channel Ca^2+^ currents of the dOrai-P288L channel could be significantly inhibited by Gd^3+^ (**Fig. 1e**). Taken together, these results show that the dOrai-P288L channel is fully active and recapitulates the properties of the STIM-activated Orai channel.

**Figure 1.**
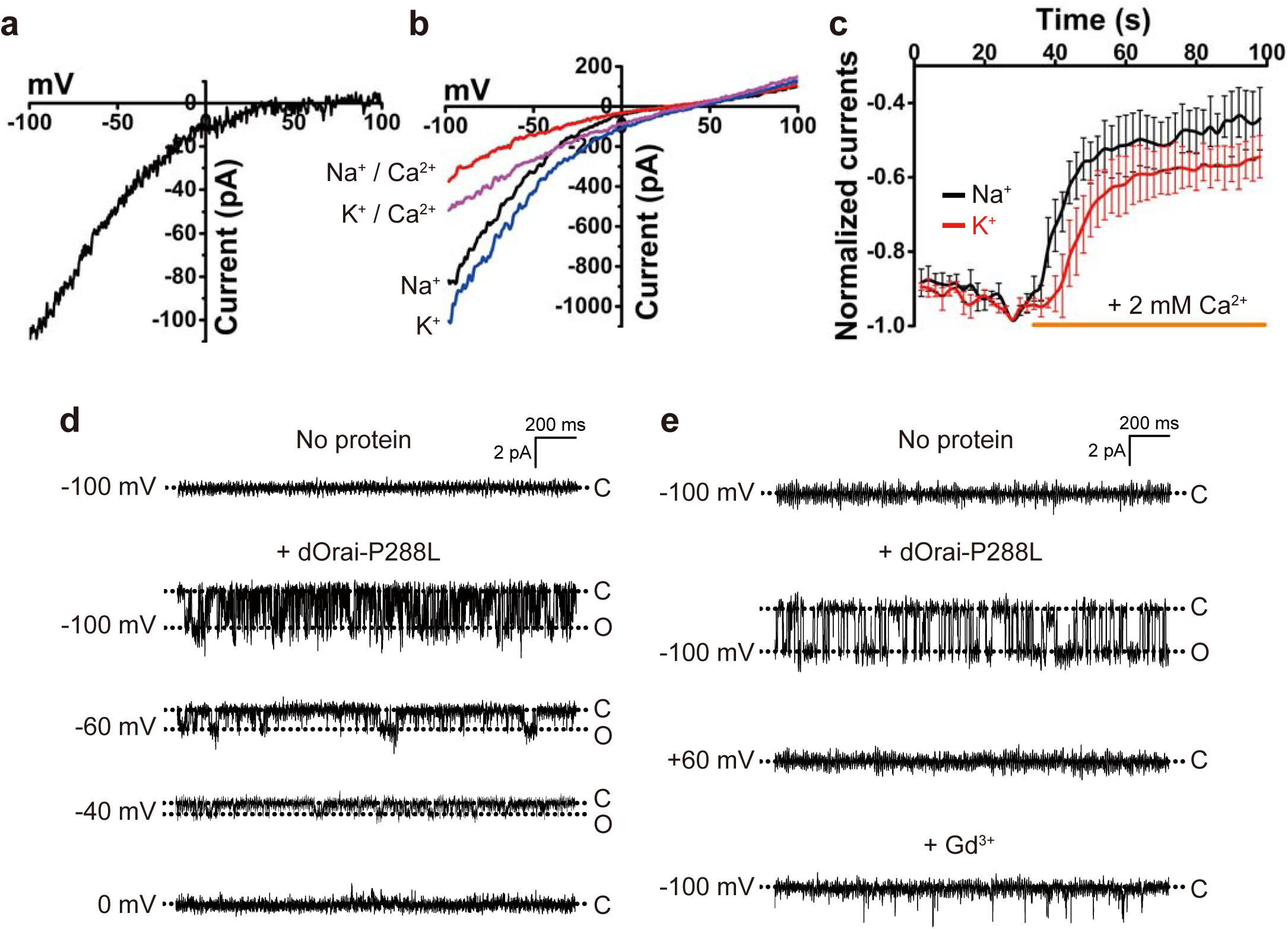
The dOrai-P288L channel is constitutively active. **a,** Current-potential (I-V) trace of the whole-cell Ca^2+^ current induced by the ramp potential. At least 3 independent experiments were conducted. **b,** Representative I-V curves of the whole-cell Na^+^ and K^+^ currents induced by the ramp potential with external Na^+^ and K^+^ recording solutions before and after the addition of 2 mM Ca^2+^. **c,** Representative time course of the normalized Na^+^ and K^+^ currents without or with the addition of 2 mM Ca^2+^. Data are shown as the mean ± S.E.M. (n=4 independent experiments). **d,** Single-channel Ca^2+^ currents evoked by the indicated stimulation voltages before and after the addition of purified proteins. **e,** The effect of 20 μM Gd^3+^ on the single-channel Ca^2+^ current.

### Overall structure of the open channel

The crystal structure of the dOrai-P288L channel was determined at the resolution of 4.5 Å by molecular replacement using the closed Orai structure (RCSB code: 4HKR) as a model. The overall architecture of the open channel shows a hexameric assembly (**Figs. 2a, 2b**). Each protomer consists of four transmembrane helices (TM1-TM4, **Fig. 2c**). Six protomers adopt a six-fold noncrystallographically symmetric arrangement, resulting in the six TM1 helices forming the ion-conducting pore in the center. The TM2 and TM3 helices from each protomer surround the six TM1 helices, forming a fence to fix the TM1 helix positions. Remarkably, each TM4 helix forms an extra-long helix, pointing into the distance on the cytosolic side. There are two hexamers in one asymmetric unit of the dOrai-P288L crystal, bound together through coiled coil interactions between the TM4 helices (Supplementary Fig. 3).

**Figure 2.**
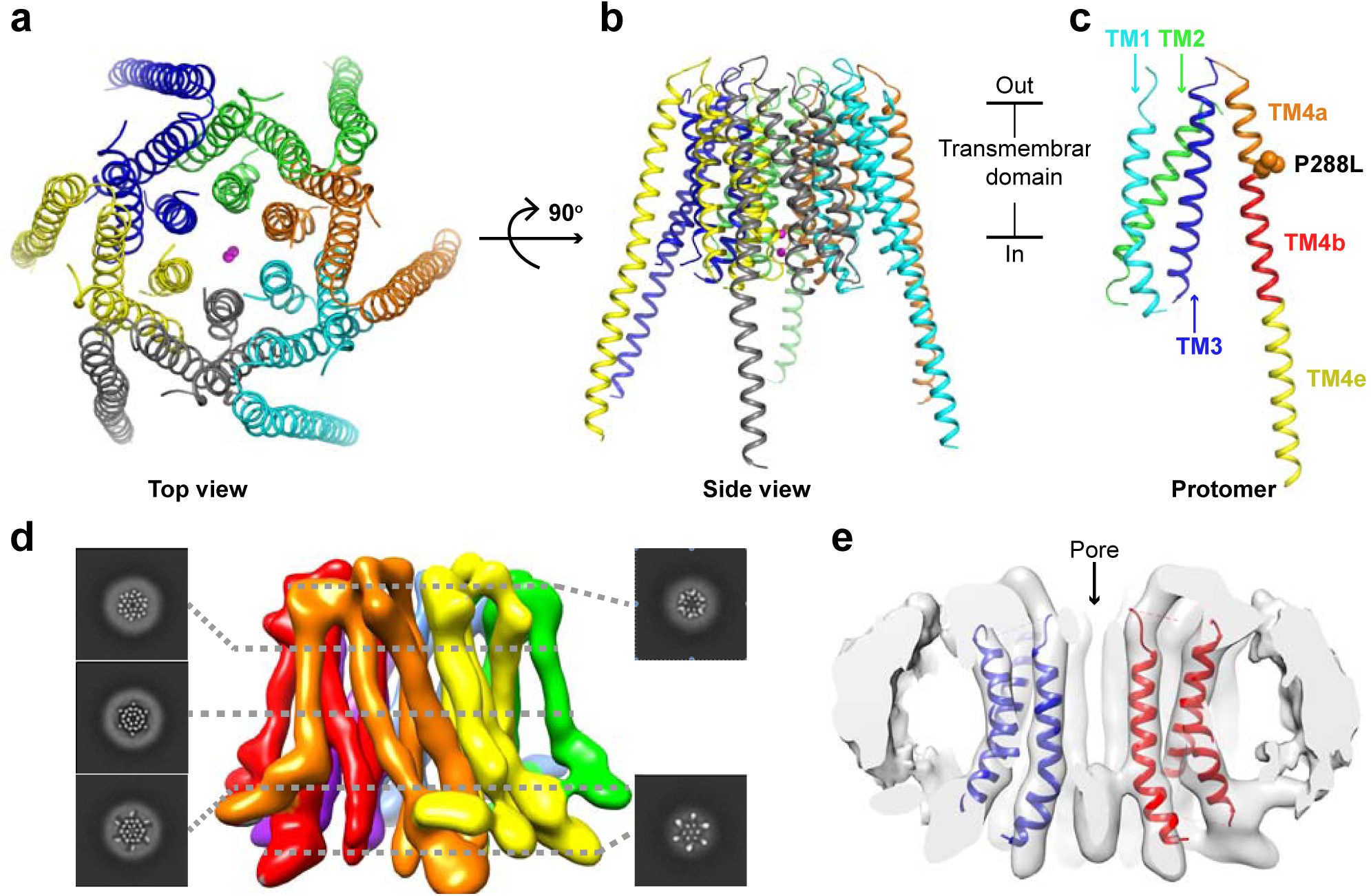
Overall structure of the open state of the Orai channel. **a,** Top view of the structure of the open Orai channel. Six protomers are colored green, blue, yellow, gray, cyan and orange, respectively. Two ions are shown as spheres in the center and are colored magenta. **b,** Side view of the structure of the open Orai channel. **c,** The architecture of one protomer. The TM1, TM2 and TM3 helices are colored cyan, green and blue, respectively. The TM4 helices are further divided into three sections, labeled TM4a (orange), TM4b (red) and TM4e (yellow). The side chains of the mutated residue P288L are shown as orange spheres. **d,** Side view of the final 3D reconstruction of the open Orai channel with slices of indicated levels. Each protomer is colored individually. **e,** Overlay of cryo-EM map (white surface) with crystal structure model (color cartoon) of the open Orai channel. Two protomers are colored blue and red, respectively.

To further confirm the open-state conformation of the dOrai-P288L channel, we took advantage of cryo-electron microscopy (cryo-EM) methodology. Negative-stain electron microscopy was used to screen different conditions including amphipols (A8-35), detergents (DDM), and reconstitution into nanodiscs. The negatively stained dOrai-P288L channels in the nanodiscs appeared monodisperse and were further subjected to cryo analysis. The samples showed orientation bias with more than 95% oriented to the same top or bottom view. We overcame this problem by using the detergent glyco-diosgenin (GDN). Because the dOrai-P288L channel was wrapped in a thick layer of detergents, the final structure was determined at the overall resolution of 5.7 Å by single-particle cryo-EM (Supplementary Figs. 4, 5). Each helix is well resolved in the final density (**Fig. 2d**). The overall cryo-EM structure of the dOrai-P288L channel is similar to its crystal structure counterpart. The crystallographic dimer across the central pore fits well into the cross-section of the cryo-EM density (**Fig. 2e**).

### Activation of the Orai channel

Structural comparison of the open and closed Orai structures revealed that the TM1-TM3 regions have similar architecture, while the TM4 helices are completely different (**Fig. 3a**). From the top view, all six TM4 helices are fully extended, and clockwise rotation occurred during opening, causing the N-terminal regions of the innermost six TM1 helices to twist outward in a counterclockwise direction. These conformational changes are also observed in the cryo-EM density of the dOrai-P288L channel (**Fig. 3b**). In contrast, the closed conformation of the Orai channel does not fit into our cryo-EM density (Supplementary Fig. 6). Surprisingly, the twisted region starts from the positively charged residue K159, which corresponds to the residue K87 in human Orai1, and proceeds to the N-terminus of the TM1 helix (**Fig. 3c** and Supplementary Fig. 7). We were not able to distinguish whether the remaining region of the TM1 helix undergoes a rotation operation ^23^ during opening in either our crystallographic structure or the cryo-EM density.

**Figure 3.**
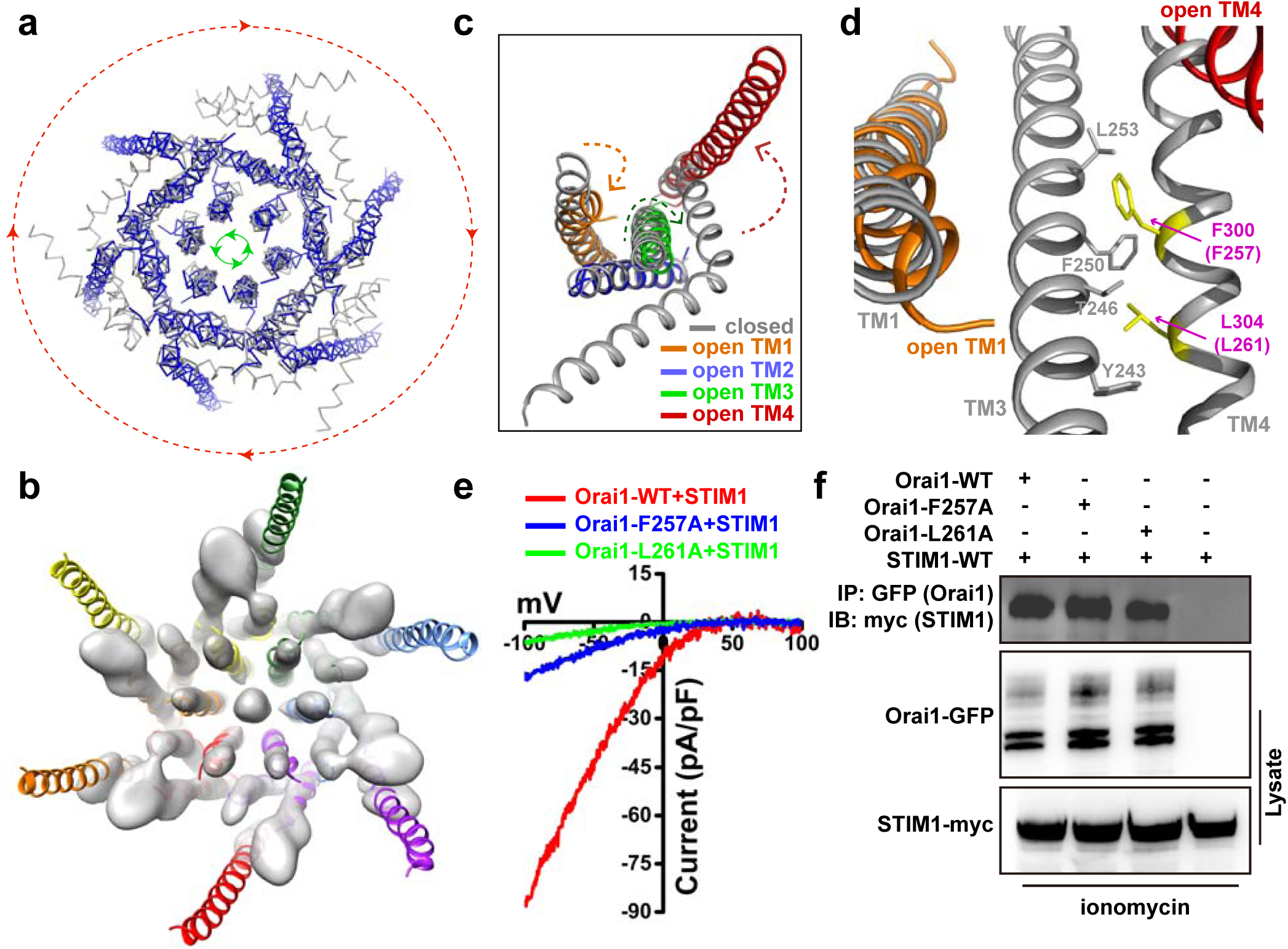
Conformation transduction pathway during channel opening. **a,** Top view of the overlay of closed (gray) and open (blue) Orai channels. During opening, the six peripheral TM4 helices rotate clockwise (red arrow direction), while the six innermost TM1 helices on the cytosolic side rotate counterclockwise (green arrow direction). **b,** Bottom view of the overlay of the open Orai channel between the crystal structure (color cartoon) and the cryo-EM map (white surface). The density in the middle presumably represents anions. **c,** Overlay of one protomer between the closed (gray) and open (color) state of the Orai channel. Arrows denote the possible helix movement during opening. **d,** Interactions between TM3 and TM4 in the closed state. Two residues, F300 and L304, are colored yellow. Residue number in human Orai1 is shown in parenthesis. **e,** Representative I-V curves of the whole-cell Ca^2+^ currents of STIM1-activated human wild-type Orai1 and mutants. **f,** Western blot analysis of coimmunoprecipitated human Orai1-GFP (wild type and mutants) with STIM1-myc.

Within each protomer, we observed a conformational transduction pathway from the peripheral TM4 helix through the middle TM3 helix to the basic section of the innermost TM1 helix (**Fig. 3c**). To confirm that such conformational transduction occurs during opening, we made two mutants (Orai1-F257A and Orai1-L261A) of wild-type human Orai1. Two residues, F257 and L261, in human Orai1 correspond to F300 and L304 in dOrai, respectively, which form hydrophobic interactions between the TM3 helix and TM4 helix in the closed conformation of dOrai (**Fig. 3d**). Upon opening, the drastic swing of the TM4b portion presumably causes the movement of the TM3 helix (**Fig. 3c**). As we expected, whole-cell patch clamp measurement showed that the Ca^2+^ currents of Orai1-F257A and Orai1-L261A after activation by STIM1 were significantly lower than those of wild-type (**Fig. 3e** and Supplementary Fig. 8a). We also used a Ca^2+^ influx assay to cross-validate this result. Two mutants (Orai1-F257A and Orai1-L261A) completely lost the extracellular Ca^2+^ influx after ER was depleted (Supplementary Fig. 8b). To ensure that the reduced channel activity was indeed caused by the reduction in channel activation, rather than an effect on attenuated STIM1-Orai1 binding, we performed co-immunoprecipitation (co-IP) and intracellular fluorescence resonance energy transfer (FRET) experiments to verify the interaction between the Orai1 mutations and STIM1. When coexpressed with STIM1-myc, wild-type and mutant Orai1-GFP were similarly able to pull down STIM1 after ionomycin treatment (**Fig. 3f**). Additionally, when YFP-tagged STIM1 and CFP-tagged wild-type and mutant Orai1 were cotransfected into HEK-293T cells, the measured apparent FRET efficiency (Eapp) values after the cells were treated with thapsigargin were similar (Supplementary Fig. 8c). These results indicate that after interference with the interaction between the TM3 helix and the TM4 helix, Orai1 cannot be activated by STIM1 but is able to associate normally with STIM1. Furthermore, in the closed conformation of dOrai1, the interaction between the basic section of the TM1 helix and the TM3 helix is mediated by three residues, L153, S154 and K157, which correspond to L81, S82 and K85 in human Orai1, respectively (Supplementary Fig. 9). Earlier reports showed that the Orai1-L81A mutant completely blocked channel function without altering the STIM1-Orai1 association, and further, the triple L81A-S82A-K85E mutant or the double L81A-S82A mutant prevented extracellular Ca^2+^ influx in the constitutively active Orai1-ANSGA mutant channel ^27^. The single replacement K85E in Orai1 resulted in a complete absence of STIM-dependent current in cells in response to Ca^2+^ store depletion ^28^. Taken together, these evidences show that the conformational transduction pathway (T4b helix --> T3 helix --> T1 helix basic section) identified in our open-state structure is critical for Orai channel activation.

### Mechanism of Ca^2+^ permeation

In the closed state of the dOrai structure, the basic section of the TM1 helix was identified and proposed to provide electrostatic repulsion in blocking cation transport ^17, 29^. During Ca^2+^ permeation, this positively charged region must be neutralized or shielded to avoid electrostatic repulsion. If this argument were true, mutations in the basic section of the TM1 helix would significantly enhance Ca^2+^ permeation through the open Orai channel. However, several published results have shown that the basic section of the TM1 helix is indispensable for Ca^2+^ permeation in Orai channel ^25, 27, 30^. In our open-state Orai structure, the electron density peaks corresponding to anions can be found around the basic section of the TM1 helix in both the crystallographic structure (**Fig. 4a**) and the cryo-EM model (**Fig. 3b**). A similar anion binding site was also reported in the crystal structure of the closed state of dOrai ^17^. These results suggest that the binding of these anions to the basic section of the TM1 helix may be necessary for Ca^2+^ permeation during channel opening. With this possibility in mind, mutating these basic amino acids will reduce the binding of anions, leading to an attenuated Orai channel and thus reconciling the above functional data. In contrast, introducing more anions on the cytosolic side of the membrane may potentiate Ca^2+^ permeation through activated Orai channels.

**Figure 4.**
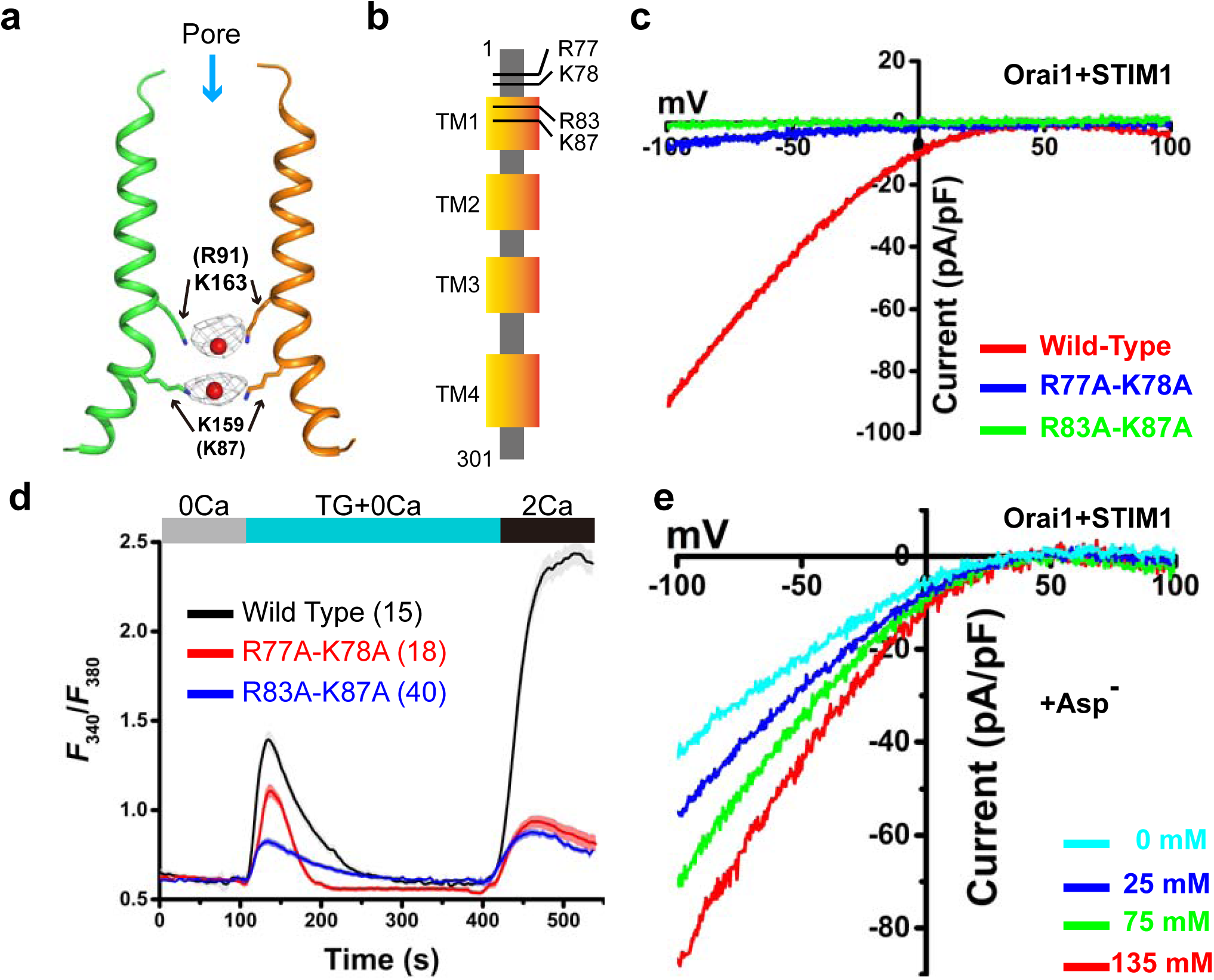
Anions facilitate Ca^2+^ permeation. **a,** Anion binding in the pore. The 2***F***o-***F***c Fourier electron density map contoured at 2.0 σ for anions is shown as a gray mesh. TM1 helices from two opposite subunits are depicted. Residues K163 and K159 are shown as sticks, and the anions are shown as red spheres. **b,** Schematic model of the human Orai1 channel. The positions of four basic residues (R77, K78, R83 and K87) are indicated. **c,** Representative I-V curves of the whole-cell Ca^2+^ currents of STIM1-activated human wild-type Orai1 and mutants. **d,** Extracellular Ca^2+^ influx in HEK-293T cells coexpressing STIM1-YFP and wild-type or mutant Orai1-GFP. **e,** Representative I-V curves of the whole-cell Ca^2+^ currents of STIM1-activated human wild-type Orai1 at the indicated concentration of sodium aspartate.

Indeed, two mutants (Orai1-R83A-K87A and Orai1-R77A-K78A) showed severely attenuated Ca^2+^ currents in the STIM1-activated Orai1 channel by whole-cell patch clamp measurement (**Figs. 4b, 4c** and Supplementary Fig. 10a) and significantly decreased store-depleted extracellular Ca^2+^ influx (**Fig. 4d**). The reduced channel activity was not caused by attenuated STIM1-Orai1 binding, because the Orai1 mutants pulled down STIM1 at a similar level to wild-type Orai after ionomycin treatment in a coimmunoprecipitation experiment (Supplementary Fig. 10b). These results reinforce the argument that the basic amino acids in this region are important for Ca^2+^ permeation in the Orai channel. More importantly, we added a certain amount of aspartate in the pipette solution to record the Ca^2+^ currents of the STIM1-activated Orai1 channel. As expected, aspartate concentration-dependent Ca^2+^ currents were observed (**Fig. 4e** and Supplementary Fig. 10c). These results suggest that the basic section of the TM1 helix may aggregate negative charges to facilitate Ca^2+^ permeation during the opening of the Orai channel.

## Discussion

In this study, we determined the structure of the constitutively active dOrai-P288L channel mutant by X-ray crystallography and cryo-EM reconstruction. The open state of the Orai channel showed a hexameric assembly. The structural comparison between the open state and the closed state reveals a conformational transduction pathway from the peripheral TM4 helix to the innermost TM1 helix. Mutations that interfere with the pathway dramatically attenuate the STIM1-activated Orai function, as demonstrated by electrophysiology. The twist of the basic section of the TM1 helix to face the cytosolic side may accommodate more anions to facilitate Ca^2+^ permeation. Therefore, we propose a model of Ca^2+^ permeation in the Orai channel (**Fig. 5**). In the closed state of the channel, the latched TM4 helix shrinks the pore on the cytosolic side. Positive charge repulsion and anion plugs block Ca^2+^ permeation. Upon opening, the TM4 helix swing twists the basic section outward to accommodate more anions. These anions not only neutralize the positive charges to reduce charge repulsion but also increase the potential gradient across the membrane, thus facilitating Ca^2+^ permeation.

**Figure 5.**
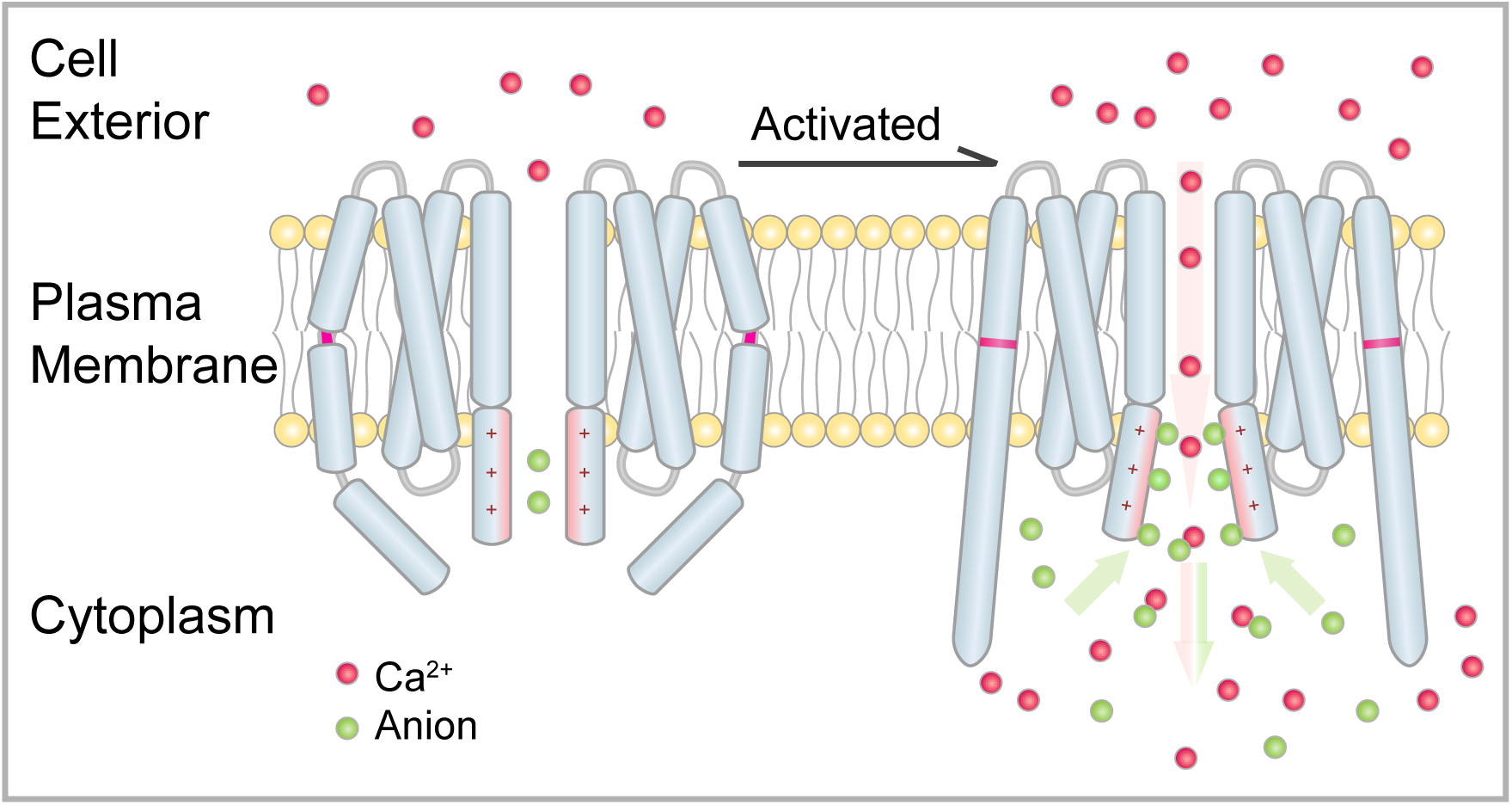
Proposed mechanism of Orai channel activation and Ca^2+^ permeation. Two opposite subunits of the Orai channel are shown. Ca^2+^ and anions are drawn as red and green spheres, respectively. In the closed state, the latched TM4 helix limits the opening of the pore on the cytosolic side. Upon activation, the swing of the TM4 helix induces the outward twist of the pore on the cytosolic side, thus aggregating anions. These anions may reduce positive charge repulsion and increase the potential gradient across the membrane, allowing Ca^2+^ permeation.

This model is consistent with many published functional studies. Zhou et al identified a “nexus” site (amino acids 261-265) within the Orai1 channel that is proposed to connect the peripheral C-terminal STIM1-binding site to the Orai1 pore helices ^27^. The structural comparison of the Orai channel structures between our open state and the published closed state clearly observed the conformational transduction pathway (T4b helix --> T3 helix --> T1 helix basic section), providing further evidence that the “nexus” site is most likely the trigger for channel activation. Furthermore, cholesterol was reported to interact with Orai1 channel and inhibit its activity through residues Orai1 L74 and Orai1 Y80 ^31^. These two residues are located within the interface between the TM1 helix and the TM3 helix. The cholesterol binding presumably interrupts the conformational transduction pathway, which explains the result that cholesterol did not affect the binding of STIM1 to Orai1 channel but attenuate the Orai1 activation ^31^.

How Orai channel conducts Ca^2+^ is a puzzling question. The Orai pore consists of an extracellular mouth, the selectivity filter, an unusually long hydrophobic cavity and an intracellular basic region. The Orai closed-state structure revealed that the narrowest region of the pore has a diameter of 6.0 Å, wide enough for a dehydrated Ca^2+^ passing through. It seems unnecessary to rotate the pore helix for channel activation as proposed by Yamashita et al ^23^. In our open state structure of the Orai channel, we did not observe the rotation operation of the pore helix. Recently, another crystal structure of the open state (H206A) of the Orai channel at low resolution (6.7 Å) was reported ^32^. Hou et al did not observe the pore helix rotation either. Thus, the mechanism of pore helix rotation is inconsistent with structural studies. Interestingly, Hou et al proposed another pore-dilation model based on their structural findings of the dilated hydrophobic cavity and also wide open intracellular basic region ^32^. However, our structure did not observe dilated hydrophobic cavity but do see the opening of the intracellular basic region. Moreover, the pore-dilation model is inconsistent with the result that mutating the intracellular basic region of the constitutively active Orai1 abolished the channel activity ^27^. Therefore, the pore-dilation mechanism less likely account for Ca^2+^ permeation of the Orai channel. While our proposed anion-assisted Ca^2+^ permeation model is reasonable to explain these results. Furthermore, it was reported that the Orai currents were inhibited by acidic but potentiated by basic intracellular solutions in various cell types ^33^. This result is nicely fit with our model since basic intracellular solutions provide more hydroxide anions while acidic solutions provide more proton cations. Both pore helix rotation and pore dilation model are hard to explain this result. In summary, our studies provide a reasonable model to clarify the molecular details of the activation and Ca^2+^ permeation of the Orai channel.

## Acknowledgements

We are grateful to the staff at the beamline BL19U of the Shanghai Synchrotron Radiation Facility and at the beamline BL-17A at Photon Factory (Tsukuba, Japan) for excellent technical assistance during data collection; We thank Dr. Ang Li for the support of electron microscopy in Nankai University, and Dr. Shenghai Chang for cryo-EM data acquisition in Zhejiang University; We thank Mrs Nannan Xiao in Nankai University for the help in imaging studies. This work was supported by National Key Research and Development Program of China (grant 2017YFA0504801 to YS; 2017YFA0504803 and 2018YFA0507700 to XZ), National Natural Science Foundation of China (grants 31570750 YS and 31870834 to YS; grant 31870736 to XY), Ph.D Candidate Research Innovation Fund of Nankai University and the Fundamental Research Funds for the Central Universities (2018XZZX001-13) to XZ.

## Author Contributions

X.F.L. did protein purification, crystallization, cell imaging and co-IP experiments; G.W. did electrophysiology studies; X.Y., J.L. and X.Z. did cryo-EM data collection; X.Y. did structure determination and cryo-EM reconstruction; Y.Y., X.C., and R.J. did protein purification and crystallization; X.L. did FRET experiment; X.F.L., G.W., X.Y. and Y.S. analyzed the data, designed the study and wrote the paper. All authors discussed the results and commented on the manuscript.

## Methods

### Plasmid construct

The human full-length Orai1 and STIM1 were inserted into the pEGFP-N1, pEYFP-N1, pECFP-N1 and PCDNA3.1/Myc-His to generate Orai1-GFP, STIM1-YFP, Orai1-CFP and STIM1-Myc. D*rosophila melanogaster* full-length Orai (Uniprot:Q9U6B8) codons were optimized for *Homo sapiens* and synthesized in Genewiz Inc. To improve the expression and stability of target protein, we chose a fragment that contains amino acid residues 132-341 and introduced two cysteine mutations (C224S and C283T). Also, P288L mutation was introduced to activate Orai channel. This Orai segment (dOrai-P288L) containing three point mutations (C224S, C283T and P288L) was cloned into pEG-BacMam vector followed by a PreScission protease cleavage site and an enhanced green fluorescent protein (eGFP) at C-terminus. All mutations were introduced using the Quick-Change Lightning Site-Directed Mutagenesis Kit (Agilent).

### Protein expression and purification

The dOrai1-P288L construct was transformed into DH10Bac *Escherichia coli* cells to generate the Bacmid. The bacmid was transfected into Sf9 cells to generate P1 baculovirus that were harvested after 72h. P1 baculovirus were amplified in Sf9 cells to generate P2 baculovirus. P2 baculovirus were added (1%-3% v/v) to HEK293S-GnTi^-^ cells when the cells reached to a density of 1.5- 2.0×10^6^ cells / mL. To increase the expression level, 10 mM sodium butyrate was supplemented and culture temperature was turned down to 30 °C for 12 h post-transduction, then further cultured for 60 h before harvesting. One liter cells were resuspended and lysed by sonicator in 100 mL buffer A (20 mM Tris-HCl pH 8.0, 200 mM NaCl and 1 mM PMSF). The membrane fraction was collected by centrifugation at 180,000 ×g in 4 °C for 1 h. The membrane was then solubilized in 70 mL buffer A plus 1% n-dodecyl-β-d-maltopyranoside (DDM), 0.2% cholesteryl hemisuccinate (CHS, Sigma) and 1× protease inhibitor cocktail (Roche) with gentle shaking in 4 °C for 1 h. Solubilized membranes were cleared by centrifugation at 100,000 ×g for 30 min. Afterwards, supernatant was mixed with 3 ml Ni-NTA beads (GE Healthcare) and incubated for 1 h. The beads were collected by a gravity column and washed by buffer B (20 mM Tris-HCl pH 8.0, 200 mM NaCl, 0.03% DDM, 0.006% CHS) and buffer C (20 mM Tris-HCl pH 8.0, 800 mM NaCl, 0.03% DDM, 0.006% CHS). Then beads were mixed with 3C protease (1:50 v/v) to cleave eGFP tag overnight. The flow through was collected and beads were washed two times by buffer B. All proteins were pooled and concentrated for further purification.

For crystal screen and optimization, the protein was injected into Superdex 200 Increase 10/300 GL (GE Healthcare) equilibrated by buffer D containing 20 mM Tris-HCl pH 8.0, 80 mM NaCl, 2 mM DTT, 0.2% n-Nonyl-β-D-glucopyranoside (NG), 0.28% n-Nonyl-β-D-Maltoside (NM) and 0.1 mg/ml lipid [POPC: POPE: POPG = 3:1:1 (w/w), Avanti Polar Lipids Inc], all peak fractions were collected and concentrated to 12 mg/mL.

For EM studies, the protein was loaded into Superdex 200 Increase 10/300 GL (GE Healthcare) equilibrated by buffer E composed of 20 mM Tris-HCl pH 8.0, 150 mM NaCl, 2 mM DTT and 0.04% glyco-diosgenin (GDN), the peak was concentrated to 0.8 mg/mL for negative-stain EM and cryo-EM grid preparation.

For single channel recording experiment, the protein was further purified using by Superdex 200 Increase 10/300 GL (GE Healthcare) equilibrated with buffer F (20 mM Tris-HCl pH 8.0, 150 mM NaCl, 2 mM DTT, 0.02% DDM and 0.004% CHS). Then peak fractions were collected and concentrated.

### Crystallization and structure determination

The protein was mixed with reservoir solution at the ratio of 1:1 by sitting drop. The reservoir solution consists of 0.05 M NaCl,0.02 M MgCl_2_, 0.1 M Sodium citrate pH 6.5, 16%-20% PEG400. The crystal was grown in 3-4 days at 17 °C. Crystal was cryoprotected in a solution contained 0.05 M NaCl, 0.02 M MgCl_2_, 0.1 M Sodium citrate pH 6.5, 45% PEG400, 0.4% NG, 0.56% NM and 0.2 mg/mL lipid (POPC:POPE:POPG=3:1:1) and was flash-frozen in liquid nitrogen.

The crystal structure of dOrai-P288L channel was determined by molecular replacement, the closed state of dOrai1 model was used as a search model in PHASER ^34^. The structure refinement was performed with the PHENIX program ^35^, and reiterated model building was performed manually in COOT ^36^. The final refinement yielding Rcrystal value of 32.57% and Rfree value of 38.83%. Structure validation was performed using MOLPROBITY ^37^. Crystallographic data and refinement statistics are reported in Supplementary Tab. 1.

### Nanodisc reconstitution

Membrane scaffold protein MSP1E3D1 was expressed and purified, and dOrai1 was reconstituted into nanodisc as previously described with modifications ^38^. Briefly, lipids (POPC:POPG:POPE = 3:1:1) were dried under argon stream for 2 h and placed in the vacuum chamber overnight to remove residual chloroform. Lipids were resuspended by sonication in buffer containing 20 mM Tris-HCl pH 8.0, 150 mM NaCl, 2 mM DTT, 20 mM DDM to final lipid concentration 10 mM. dOrai1-P288L, MSP1E3D1, and lipids were mixed at a molar ratio of 1:4:140. The mixture was incubated on ice for 1 h. Afterwards, Bio-beads (20 mg/mL, Bio-Rad) was added to the mixture and gentle shaking at 4°C to start the reconstitution. After 3 h, Bio-beads were exchanged and then rotated overnight. The supernatant was loaded into Superdex 200 Increase 10/300 GL equilibrated with Gel-filtration buffer (20 mM Tris-HCl pH 8.0, 150 mM NaCl, 2 mM DTT), and the peak fraction was collected and concentrated to 0.8 mg/mL for negative-stain EM and cryo-EM grid preparation.

### EM grid preparation and data collection

For cryo-EM sample preparation, an aliquot of 3.5 μl fresh sample of concentration 0.8 mg/ml was applied to a glow-discharged holy carbon grid (Quantifoil, R2/1, 200 mesh). The grids were blotted for 5 s under 100% humidity at 20 °C, and plunged into liquid ethane cooled by liquid nitrogen with an FEI Vitrobot Mark IV (FEI Company). The grid was loaded onto FEI Titan Krios electron microscope with a K2 Summit direct electron counting detector (Gatan). Movies were collected by *SerialEM* with the under-focus range of 1.5-2.5 μm and under a magnification of 29,000x in super resolution mode, corresponding to a pixel size of 0.507 Å on the specimen level. Each movie was dose fractionated to 40 frames with 0.2 s exposure time per frame. With a dose rate of 1.6 counts per physical pixel per second, resulting in a total dose of 49.8 e^-^/ Å^2^. A total of 5182 movies were acquired from 3 grids in an image session of 120 h.

### Cryo-EM Image processing

The record movies were processed by MotionCorr2 ^39^ for a 5 x 5 patches drift correction with dose weighting and binned 2-fold, resulting in a pixel size of 1.014 Å/pixel. The non-dose-weighted images were used for CTF estimation by CTFfind 4.18 ^40^. The dose-weighted images were used for particles picking. 191,921 particles were semi-automatically picked by Gautomatch (http://www.mrc-lmb.cam.ac.uk/kzhang/) and extracted by Relion-2.1 ^41^ in a box size of 220 pixels. 2D classification was performed in Relion-2.1 to remove contaminations, ice, and bad particles, yielding 110,752 good particles. The initial model was generated using the pdb file of the crystal structure by Molmap command in Chimera ^42^ and low passed filtered to 10 Å resolution for 3D classification in Relion-2.1. A round of 3D classification into 6 classes yielded 1 reasonable reconstruction (Class1, 22,442 particles) for further refinement. Refinement of the selected particles generated a map with an average resolution of 5.7 Å for the C3 symmetry. The map was sharpened with B-factors of −150 Å^2^. ResMap ^43^ method was used to calculate the local resolution of our final dOrai-P288L map. All the figures were prepared in PyMol software (https://pymol.org/) and Chimera ^42^. Data collection and reconstruction statistics are presented in Supplementary Tab. 2.

### Cell Culture and transfection

HEK293T was cultured in DMEM (Sigma) supplemented with 10% fetal bovine serum (FBS, PAN) at 37 °C with 5% CO_2_. *Spodoptera frugiperda (Sf9) cells* grew in Insect X-Press (Lonza) at 27 °C with. HEK293S GnTi^-^ was cultured in Freestyle 293 (Invitrogen) at 37 °C with 5% CO_2_. Bacmid was transfected into *Spodoptera frugiperda (Sf9) cells* using X-tremeGENE HP DNA Transfection Reagent (Roche). All plasmids were transfected into HEK293T cells using Fugene6 (Promega).

### Whole-cell Patch clamp recordings

HEK293T cells were transiently transfected with either GFP tagged dOrai-P288L alone or human Orai1-GFP (wild type and mutants) plus human STIM1-YFP for 12–24 h at 37 °C with 5% CO_2_. Transfected cells were then digested by trypsin to be plated onto 35 mm dishes for cultivation at least 3 h before electrophysiology. The glass pipettes were pulled to a suitable shape through using a P-97 glass microelectrode puller (Sutter Instrument) and polished with an MF-830 (Narishige). For recording dOrai-P288L currents, the internal solution was 160 mM NMDG, 10 mM HEPES and 10 mM glucose (pH adjusted to 7.2 with HCl). The resistance was 3-5 MΩ after filled with the internal recording solution. For recording Ca^2+^ currents, the external Ca^2+^ solution was 105 mM CaCl2, 10 mM HEPES and 10 mM glucose (pH adjusted to 7.4 with Trizma base). For recording Na^+^ or K^+^ currents, the external Na^+^ or K^+^ solution was 160 mM NaCl or 160 mM KCl, 10 mM HEPES and 10 mM glucose (pH adjusted to 7.4 with Trizma base). All experiments were conducted at room temperature with the stimulation voltage: 50 ms voltage step to –100 mV from a holding potential of 0 mV, followed by a voltage ramp increasing from –100 to + 100 mV in 50 ms with a frequency of 0.5 Hz. In the whole-cell configuration, the 0 Ca^2+^, 0 Na^+^ and 0 K^+^ external solutions were respectively replaced with the external Ca^2+^, Na^+^ and K^+^ solutions through a peristaltic pump to record the corresponding currents. For recording human Orai1 currents, the internal solutions are 135 mM L-Aspartic acid or 100 mM D-Sorbitol plus 75 mM L-Aspartic acid or 200 mM D-Sorbitol plus 25 mM L-Aspartic acid or 250 mM D-Sorbitol plus 0 mM L-Aspartic acid, 10 mM EGTA, 10 mM HEPES and 8 mM MgCl_2_ (pH djusted to 7.2 with CsOH). The extracellular 0 Ca2^+^ solution is 130 mM NaCl, 4.5 mM KCl, 22 mM MgCl_2_, 10 mM TEA-Cl, 10 mM D-glucose and 5 mM Na-HEPES (pH 7.4). The extracellular Ca2^+^-containing solution is 130 mM NaCl, 4.5 mM KCl, 20 mM MgCl_2_, 2 mM CaCl_2_, 10 mM TEA-Cl, 10 mM D-glucose and 5 mM Na-HEPES (pH 7.4). Whole-cell currents were amplified with an Axopatch 700B and digitized with a Digidata 1550A system (Molecular Devices). All currents were sampled at 10 kHz and low-pass filtered at 2 kHz through pCLAMP software (Molecular Devices). Origin 9.0 software (OriginLab Corp.) also was used for data analysis.

### Planar lipid bilayer formation

The planar lipid bilayer was formed on a Nanion NPC-1 chip from the giant unilamellar vesicles (GUV) generated through the electroformation methods (Nanion, Germany). Materials required for GUV formation are listed as following: DPhPC (Diphytanoylsn-glycero-3-phosphatidylcholine) lipids (Avanti Polar Lipids), cholesterol (Sigma), chloroform (Carl Roth), sorbitol (Sigma), NPC-1 chip with an aperture and Vesicle Prep Pro (with electroformation chambers containing ITO slides, O-ring). DPhPC and cholesterol were dissolved in chloroform to 10 mM and 1 mM respectively, and sorbitol was dissolved in double distilled water to 1 M. An electroformation protocol of 3 V peak to peak and 5 Hz frequency at 37°C lasting 188 minutes was used to increase the number of GUV. GUVs were verified under the microscope.

### Single channel recordings

The internal solution contained 50 mM CsCl, 10 mM NaCl, 60 mM CsF, 20 mM EGTA, and 10 mM HEPES (pH adjusted to 7.2 with CsOH; 285 mOsmol). The internal solution was firstly added onto both surfaces of the chip aperture. GUVs were then added and positioned onto the aperture by a slight negative pressure (planar lipid bilayers formation on the Port-a-Patch, Nanion). The GUV burst to form the planar lipid bilayer with a high seal resistance about tens to hundreds of GΩ after touching the glass chip. The enhancer solution (Nanion) containing 80 mM NaCl, 3 mM KCl, 10 mM MgCl_2_, 35 mM CaCl2 and 10 mM HEPES (pH adjusted to 7.4 with NaOH; 298 mOsmol) was used as the external recording solution. Single channel recordings were conducted using the PatchMaster software with a HEKA EPC10 USB amplifier (HEKA). Current signals were acquired using PatchMaster software with a 10-kHz sampling frequency and were filtered at 2 kHz. The holding potential was 0 mV. Stimulation potentials were given from the *cis* side of the glass chip chamber. Zero potential was assigned by convention to the *trans* side of the chip (the grounded side). Purified proteins were added to the external surface of the chip (the *trans* side). Acquired data were analyzed by using Origin 9.0. All experiments were performed at room temperature.

### Co-immunoprecipitation and Western blotting

Cells were harvested after 24 h of plasmids transfected. Before collected, cells were washed three times with TBS and treated by ionomycin for 5 min at room temperature. Cells were lysed by buffer containing 20 mM Tris-HCl pH 7.5, 150 mM NaCl, 0.5% Triton X-100, Roche protease inhibitor cocktail, 1 mM EGTA at 4 °C. After 1 h, cell supernatant was collected by centrifugation at 20,000 ×g, 4 °C, 30 min. 1 mL cell lysates were incubated with 15 μL GFP-Trap (ChromoTek) for 2 h at 4 °C. The beads was washed three or four times by cold washing buffer composed of 20 mM Tris-HCl pH 7.5, 150 mM NaCl, 0.1% Triton X-100, 1 mM EGTA, followed by heating in 30 μL buffer containing SDS gel-loading at 95 °C for elution. Afterwards, eluted protein was loaded onto 11% SDS-PAGE gel and transferred to PVDF membranes for western blotting. The primary antibody of immunoblot analysis is anti-myc (Sigma-Aldrich, M5546) and anti-GFP (Abcam, ab6556). The protein/antibody complexes were detected using chemiluminescence.

### FRET measurements

FRET was measured as described earlier ^44^. In brief, Orai1-CFP (donor) and STIM1-YFP (acceptor) were transiently transfected into HEK293T cells. Imaging was measured utilizing Leica DMI6000B microscope equipped with high-speed fluorescence-external filter wheels for a CFP-YFP index-based FRET experiment. Images were captured every 10 s using a 63× oil / 1.4 oil objective (Leica) and controlled by LAS software. Data was analyzed by the Biosensor Processing Software 2.1 package in MATLAB R2014a and ImageJ. Bleed-through and direct excitation containment were subtracted according to the formula Fc = I_DA_ - aIAA – dIDD (assuming *b* = *c* = 0), where d represents bleed-through of CFP through the FRET filter (d = 0.715) and a represents direct excitation factor of YFP through the FRET filter (a = 0.173). The apparent FRET efficiency (Eapp) was analyze using the equation Eapp = Fc/(Fc + GI_DD_), in which Eapp represents the fraction of donor (ECFP) exhibiting FRET. The D1ER cameleon was used to determine the instrument specific constant G (G = 3.458).

### Intracellular Ca^2+^ measurement

Intracellular Ca^2+^ measurement was performed on HEK-293T cells 20-24 h after transfection with STIM1-YFP and Orai1-GFP. Cells were treated with 2 μM fura-2/AM (Sigma) in standard Ringer’s solution (140 mM NaCl, 10 mM HEPES pH 7.4, 10 mM Glucose, 0.8 mM MgCl_2_, 2.8 mM KCl, 2 mM CaCl2) for 30 min. All these processes are done at room temperature. Fluorescence imaging was measured using a Leica DMI6000B microscope with a 40× oil-immersion objective lens controlled by LAS software. Consecutive excitation occurred at 340 and 380 nm, and emission was collected at 510 nm. Intracellular Ca^2+^ concentration is shown as the 340/380 ratio (F340/F380) obtained from 15-40 cells.

### Data availability

Atomic coordinates have been deposited in the Protein Data Bank under accession number 6AKI. The cryo-EM density maps of the dOrai-P288L channel have been deposited in the Electron Microscopy Data Bank under accession number EMD-9641.

---

**Supplemental Table 1.** Data collection and refinement statistics

|  |  |
| --- | --- |
| Crystal name | **dOrai-P288L** |
| Space group | ***P2*<sub>1</sub>** |
| Unit cell |  |
| ***a*, *b*, *c* (Å)** | ***a*=118.107, *b*=90.659, *c*=178.721** |
| ***α*, *β*, *γ* (°)** | ***α*=*γ*=90, *β*=106.31** |
| Wavelength (Å) | 0.9800 |
| Resolution range (Å) | 50-4.50 (4.66-4.50)** |
| No. of unique reflections | 17,659 (850)** |
| Redundancy | 3.5 (2.7)** |
| ***R*<sub>sym</sub> (%)\*** | **12.4 (58.7)\*\*** |
| ***I*/*σ*** | **7.94 (2.00)\*\*** |
| Completeness (%) | 80.7 (39.8)** |
| Figure of merit |  |
| Refinement |  |
| ***R*<sub>free</sub> (%)<sup>†</sup>** | **38.83** |
| ***R*<sub>crystal</sub> (%)<sup>‡</sup>** | **32.57** |
| **RMSD<sub>bond</sub> (Å)** | **0.004** |
| **RMSD<sub>angle</sub>(°)** | **0.93** |
| Number of |  |
| Protein atoms | 12,372 |
| Ligand atoms | 4 |
| Solvent atoms | 0 |
| Residues in (%) |  |
| Ramachandran favored | 95.08 |
| Ramachandran allowed | 4.57 |
| Ramachandran outliers | 0.17 |
| Rotamer outliers | 0 |
| Average B factor (Å<sup>2</sup>) of |  |
| Protein | 122.88 |
| Ligand | 87.74 |
| Solvent | - |

\* $R_{\text{sym}} = \sum_j |I_j - \langle I \rangle| / \sum \langle I \rangle$ where $I_j$ is the intensity of the *j*th reflection and $\langle I \rangle$ is the average intensity.
\*\* the highest resolution shell.
<sup>‡</sup> $R_{\text{free}}$ , calculated the same as $R_{\text{crystal}}$ , but from a test set containing 5% of data excluded from the refinement calculation.

**Supplemental Table 2.**
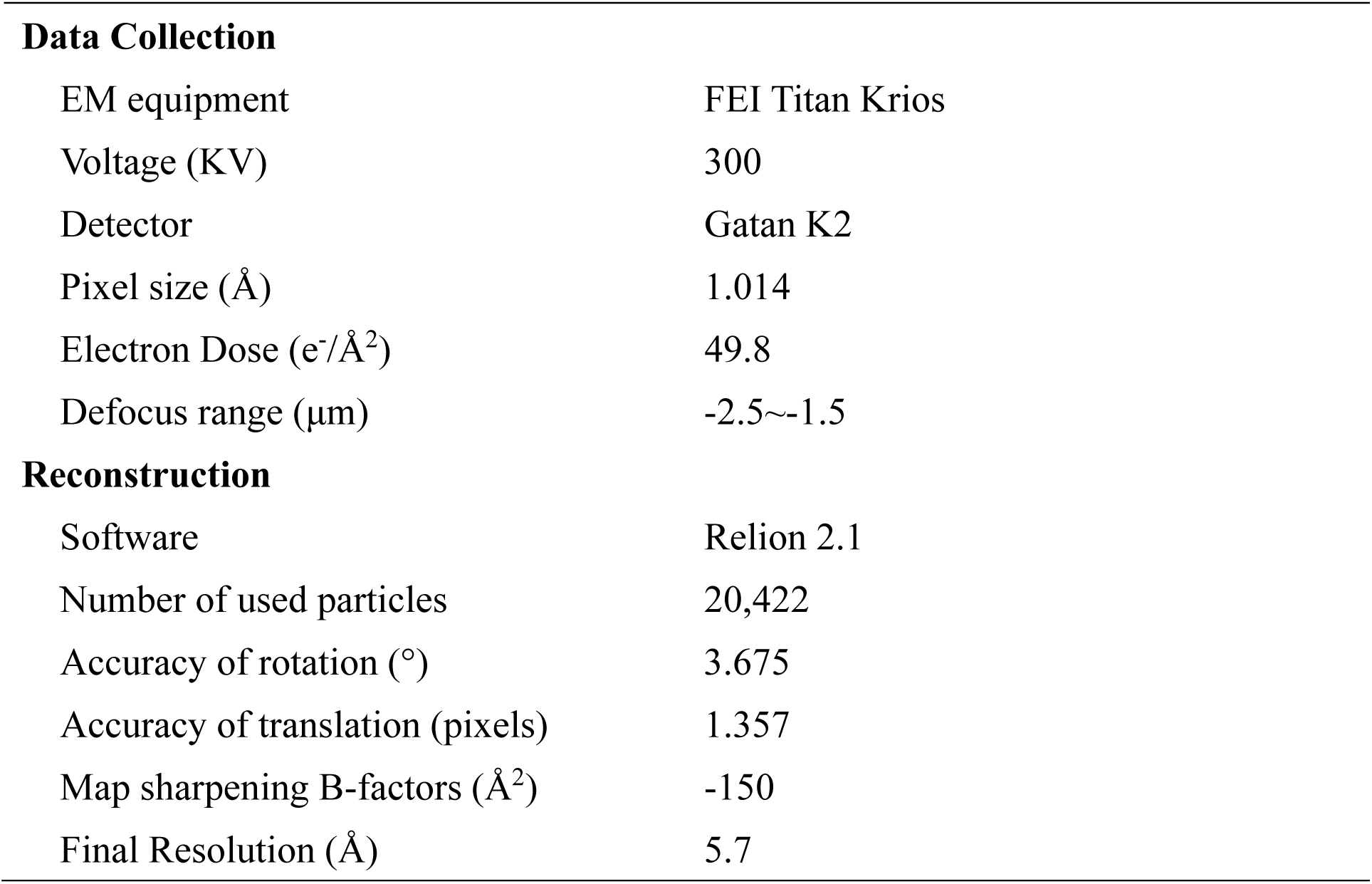
CryoEM data collection and Reconstruction statistics

**Data Collection**
|  |  |
| --- | --- |
| EM equipment | FEI Titan Krios |
| Voltage (KV) | 300 |
| Detector | Gatan K2 |
| Pixel size (Å) | 1.014 |
| Electron Dose (e<sup>-</sup>/Å<sup>2</sup>) | 49.8 |
| Defocus range (μm) | -2.5~-1.5 |
| **Reconstruction** |  |
| Software | Relion 2.1 |
| Number of used particles | 20,422 |
| Accuracy of rotation (°) | 3.675 |
| Accuracy of translation (pixels) | 1.357 |
| Map sharpening B-factors (Å<sup>2</sup>) | -150 |
| Final Resolution (Å) | 5.7 |

**Supplemental Figure 1.**
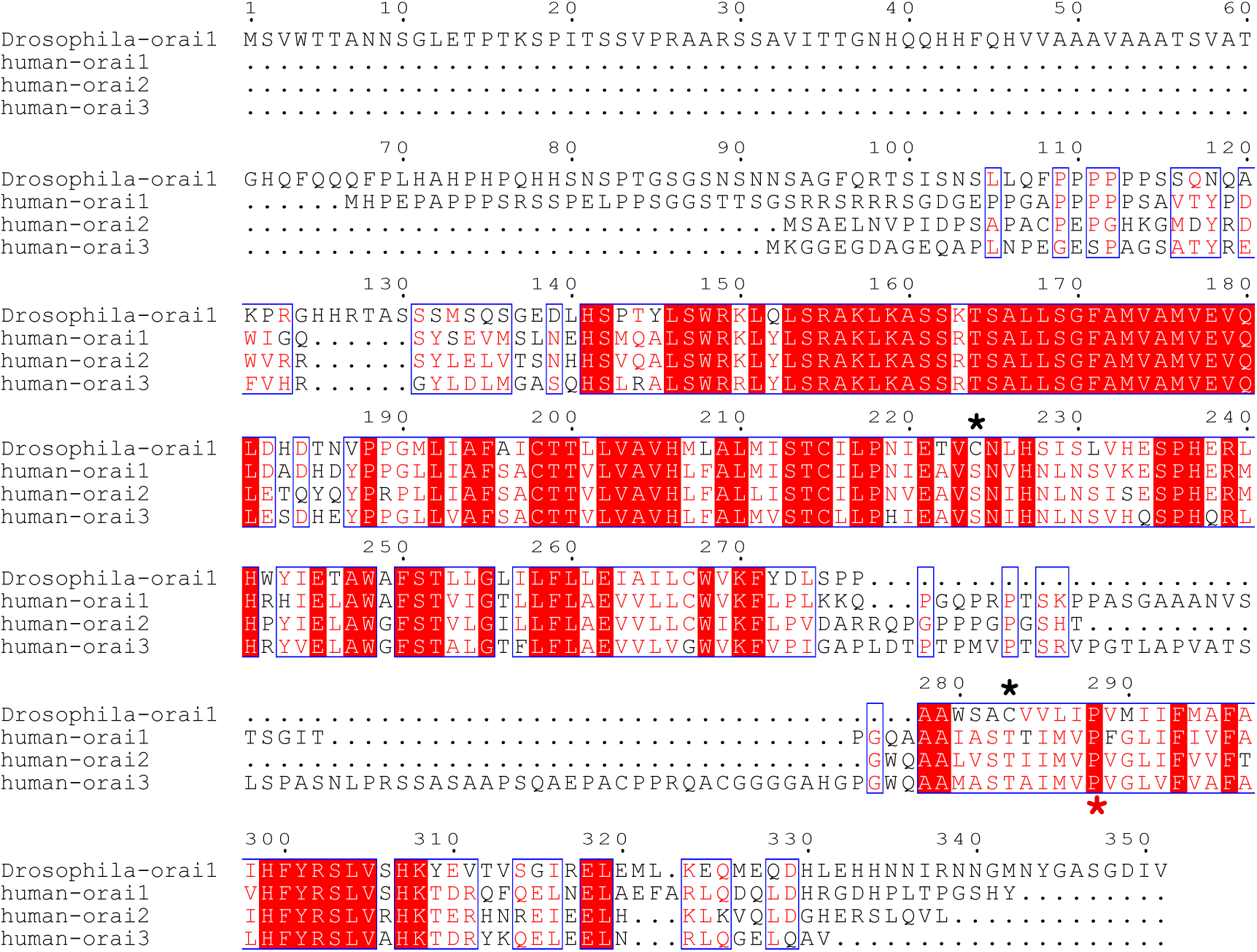
Sequence alignment. The residues that are conserved among all four proteins are highlighted in red. The accession numbers for the sequences in the alignment are Q9U6B8 for fly Orai, Q96D31 for human Orai1, Q96SN7 for human Orai2 and Q9BRQ5 for human Orai3, respectively. The star symbol denotes the amino acids mutated in structural studies.

**Supplemental Figure 2.**
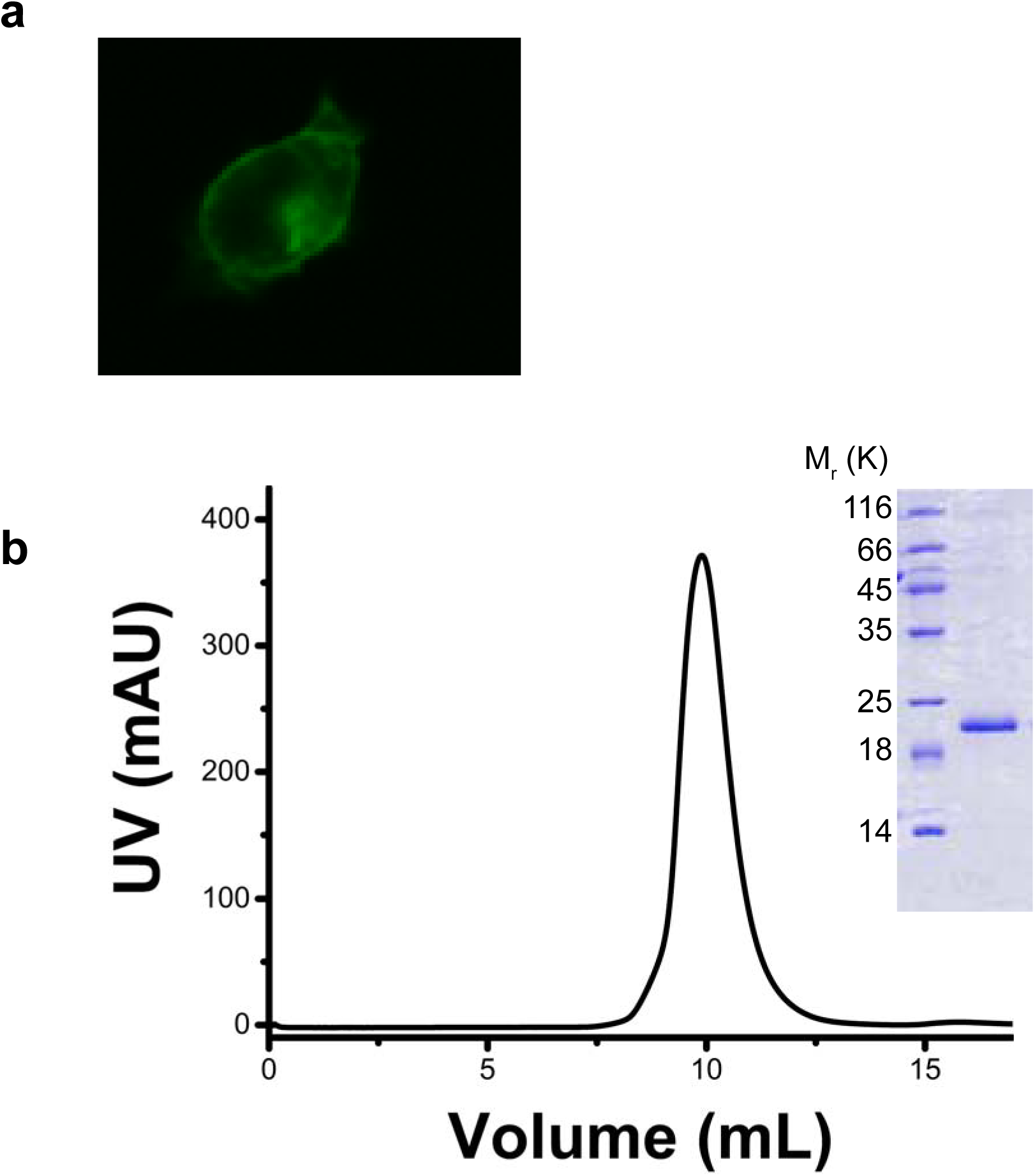
Biochemical properties of the dOrai-P288L channel. **a,** Fluorescence microscopy picture of the dOrai-P288L channel localized at the surface of HEK-293T cells. **b,** Gel-filtration profile of the purified dOrai-P288L channel. The inset shows the purity of the dOrai-P288L channel observed by SDS-PAGE.

**Supplemental Figure 3.**
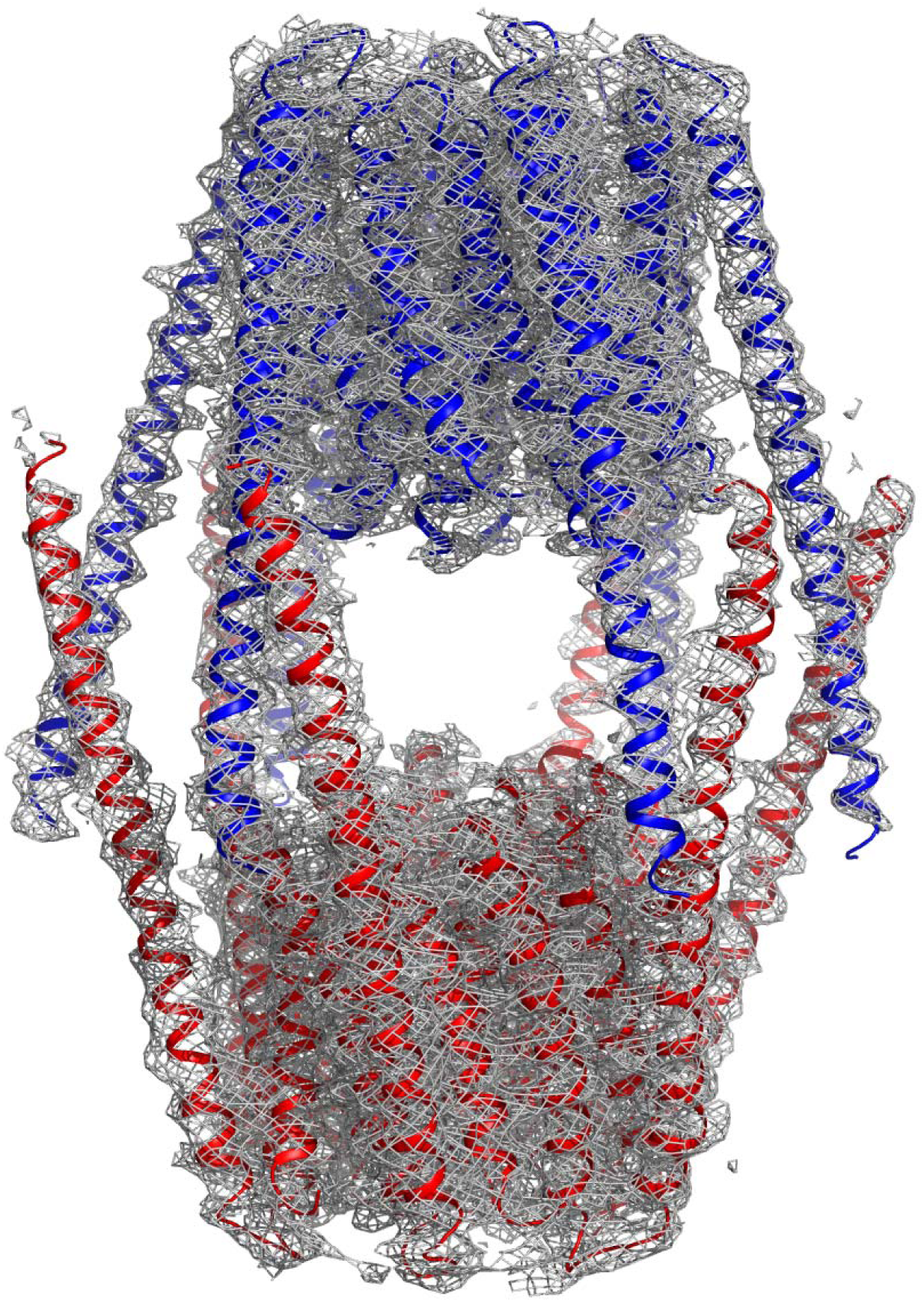
Molecule packing in one asymmetric unit of dOrai-P288L crystal. Two hexamers of the dOrai-P288L channel are shown in blue and red. The electron density is drawn in gray.

**Supplemental Figure 4.**
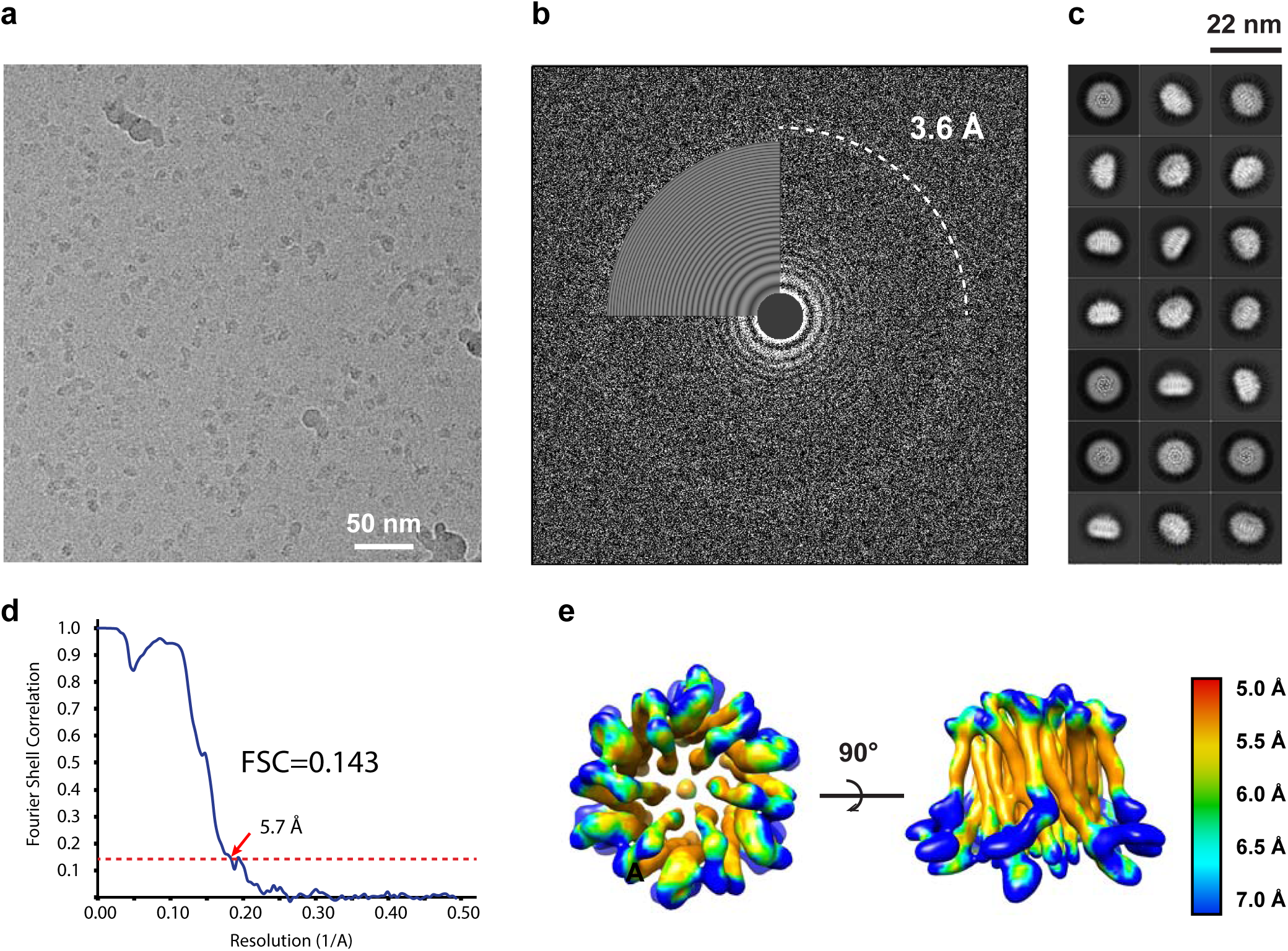
Cryo-EM structure determination and resolution assessment of the dOrai-P288L channel. **a,** A drift-corrected cryo-EM micrograph of the dOrai-P288L channel. **b,** Ctffind showed Thon rings in the Fourier spectrum of the image in panel a. **c,** Selected 2D class averages of the dOrai-P288L channel. **d,** The “gold-standard” FSC coefficient curve of the final reconstruction showed an overall resolution of 5.7 Å. **e,** Local resolution estimation by ResMap.

**Supplemental Figure 5.**
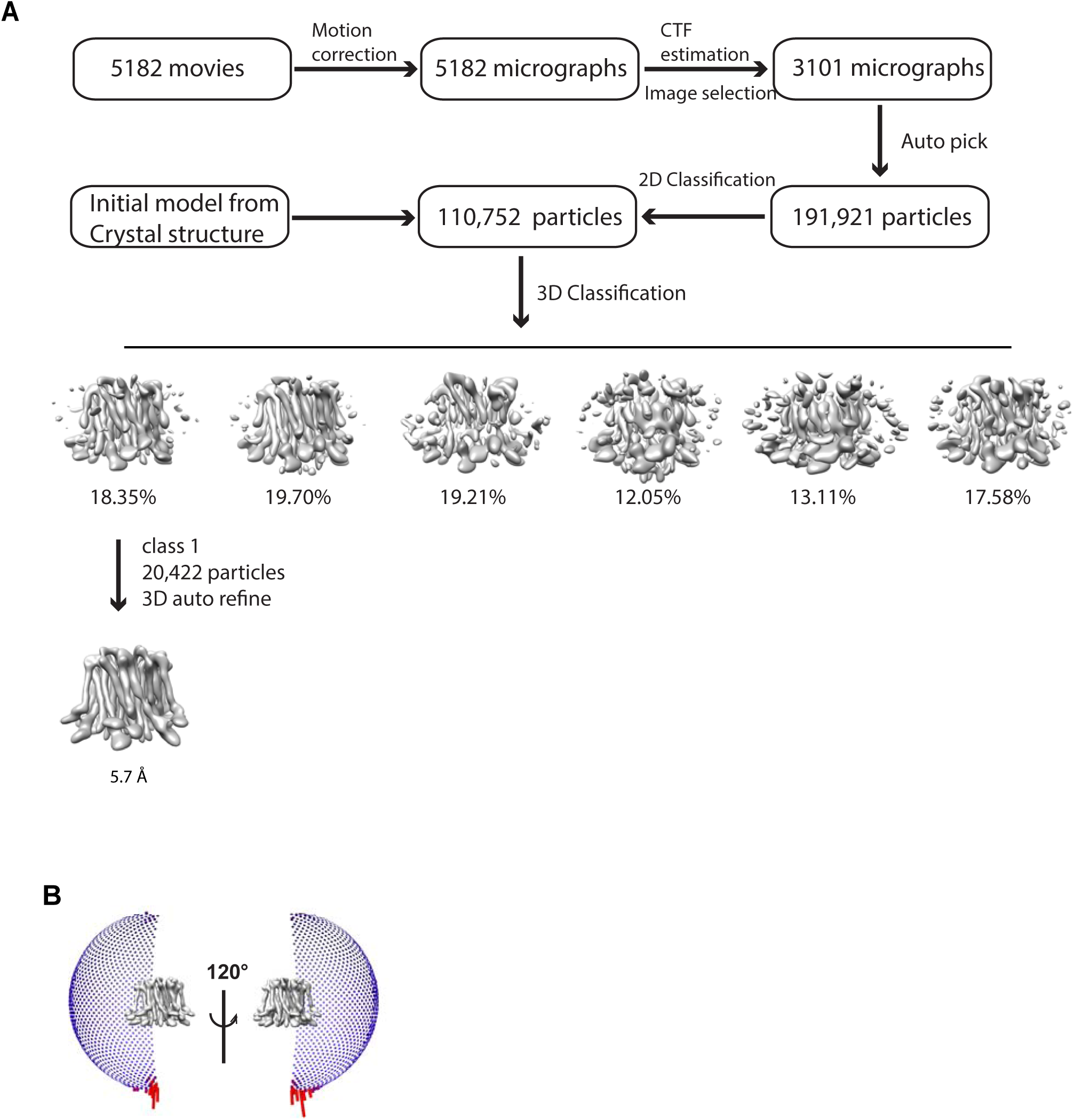
Cryo-EM data processing of dOrai-P288L channel. **a,** Flow chart of whole data processing. **b,** Orientation distribution of particles included in the final reconstruction.

**Supplemental Figure 6.**
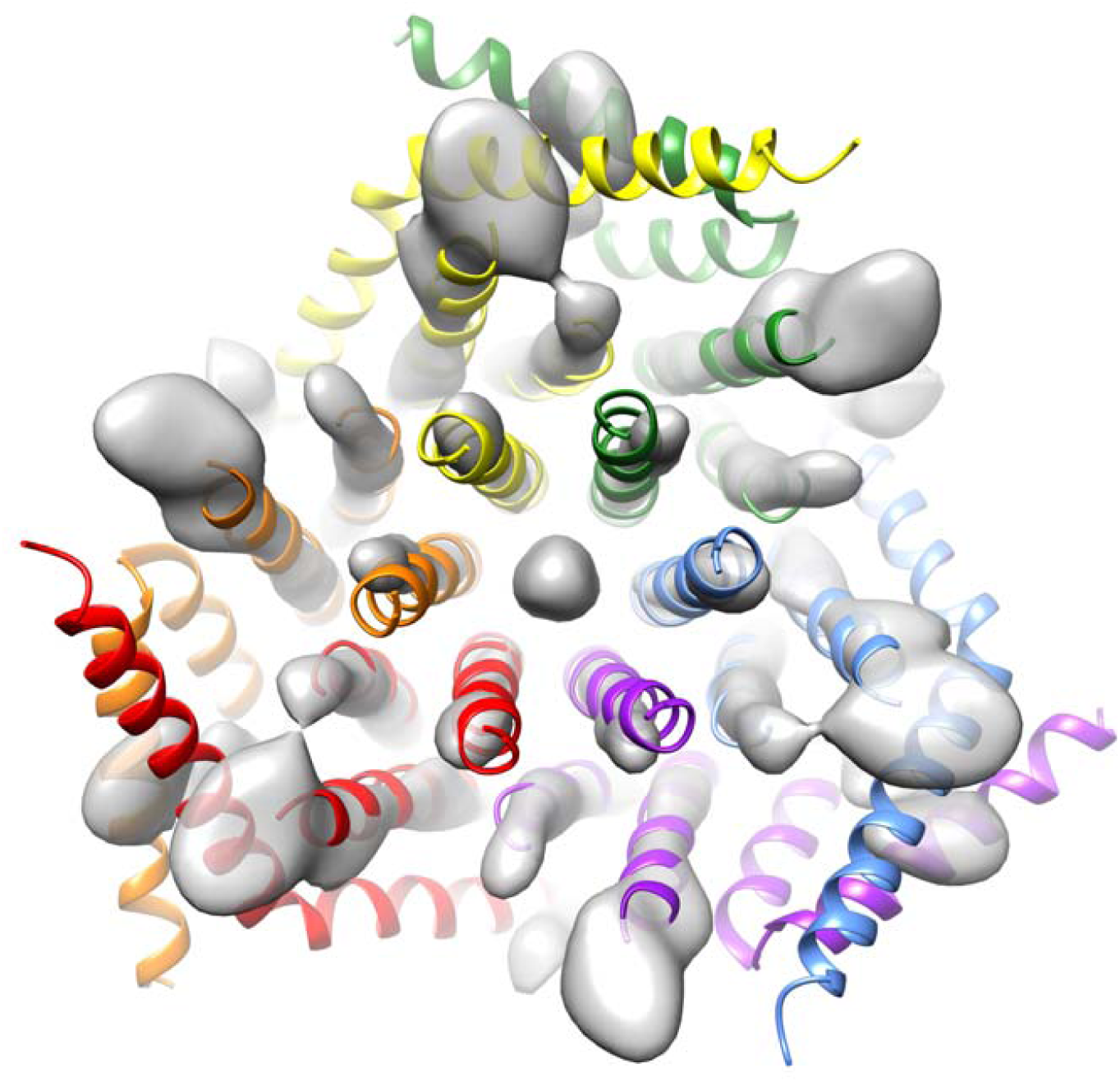
Bottom view of the overlay of the crystal structure (color cartoon) of the closed Orai channel and the cryo-EM map (white surface) of the dOrai-P288L channel. The crystal structure cannot be fitted into the cryo-EM map.

**Supplemental Figure 7.**
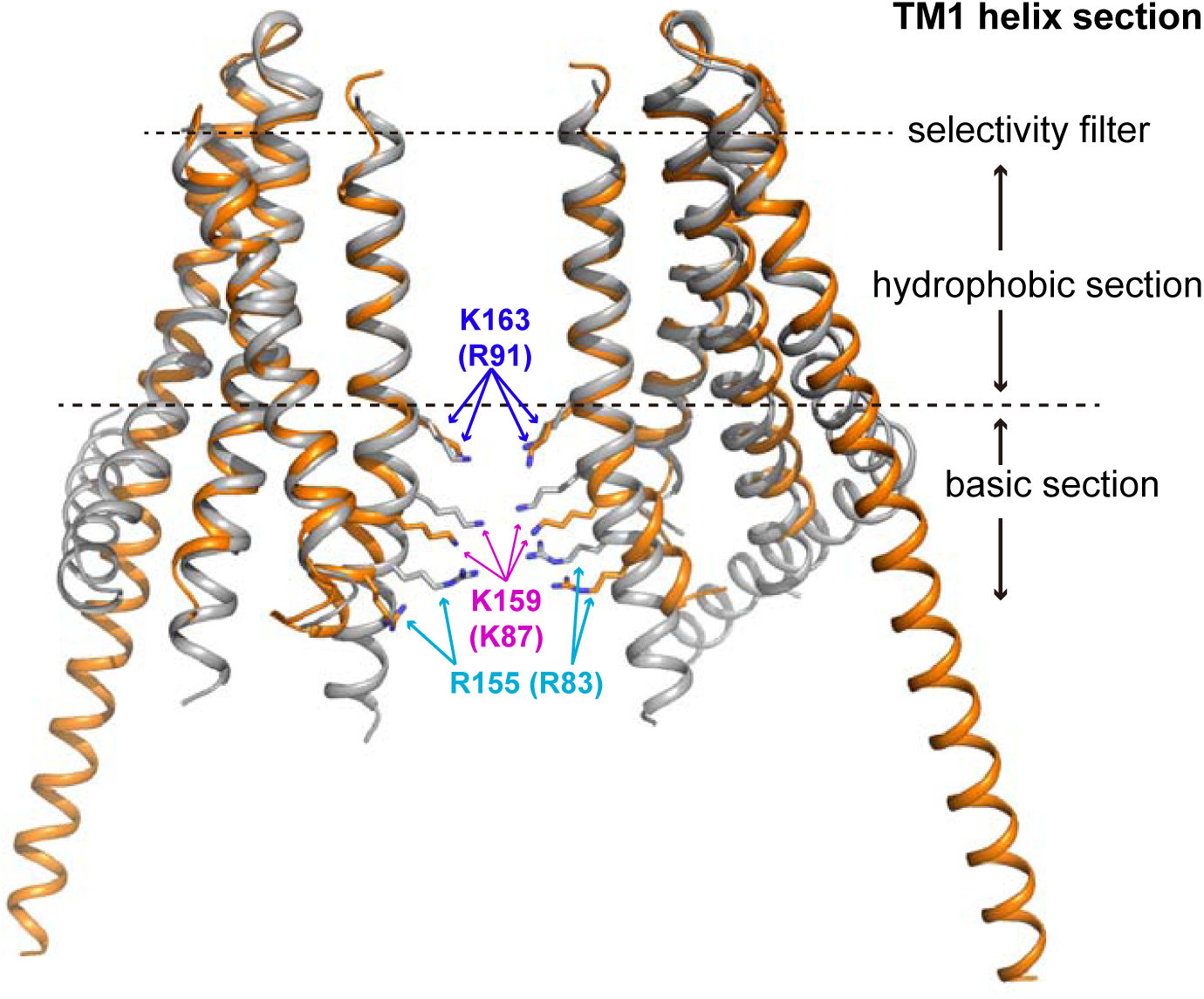
Conformational changes between the closed and open Orai channels. The closed Orai channel is colored gray, and the open Orai channel is colored orange. Two opposing protomers are shown. Side chains of residues K163, K159 and R155 are shown as stick models. The amino acid numbers of human Orai1 are shown in parentheses.

**Supplemental Figure 8.**
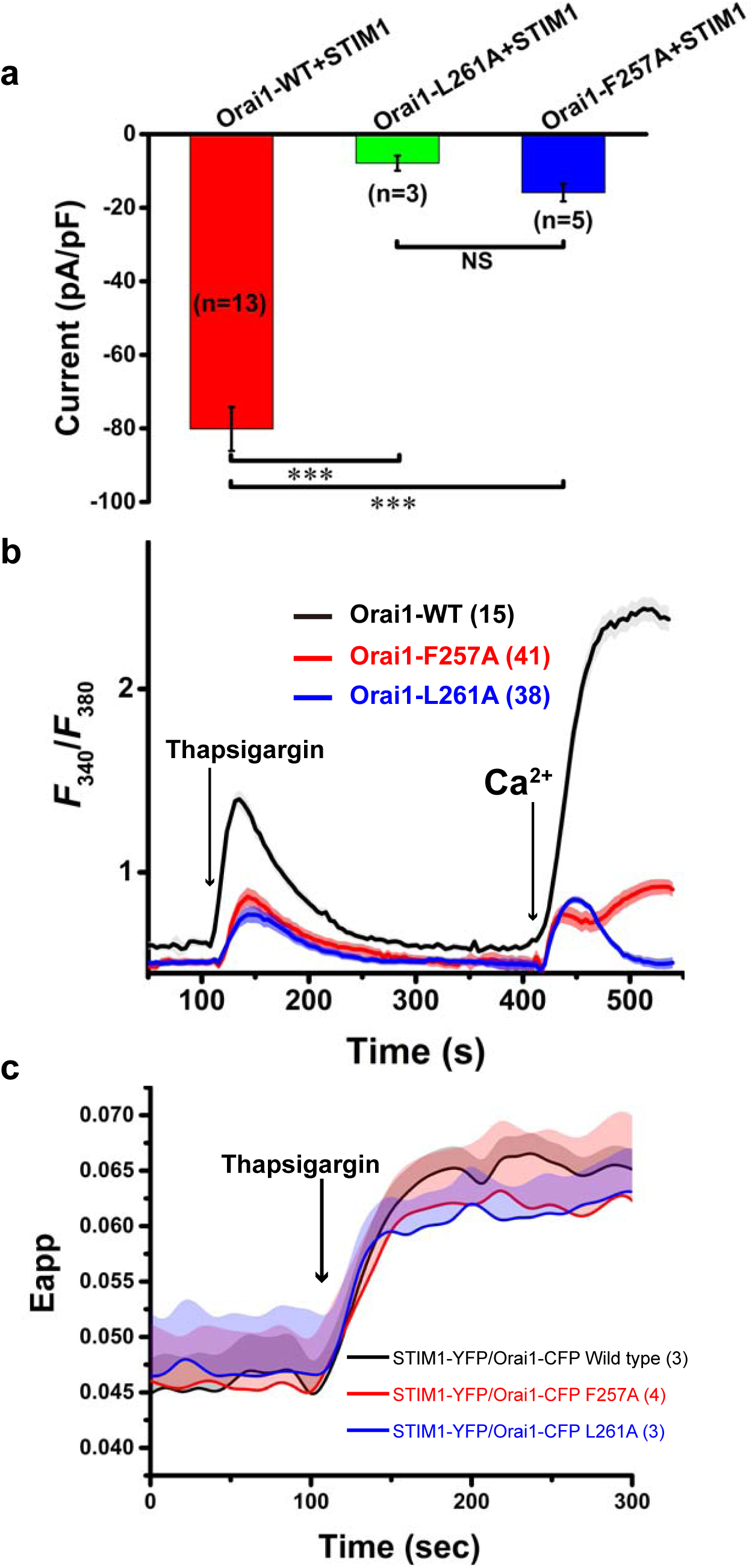
TM3-TM4 hydrophobic interaction is essential for STIM1-dependent Orai1 activation. **a,** Bar graphs of whole-cell Ca^2+^ currents of STIM1-activated human wild-type and mutant Orai1 channels (Orai1, Orai1-L261A and Orai1-F257A). **b,** Extracellular Ca^2+^ influx in HEK-293T cells coexpressing STIM1-YFP and wild-type or mutant Orai1-GFP. **c,** FRET between STIM1-YFP (acceptor) and wild-type or mutant Orai1-CFP (donor) coexpressed in HEK-293T cells. The curves of F257A and L261A are colored red and blue, respectively. The number of analyzed cells is indicated. Error bars denote S.E.M.

**Supplemental Figure 9.**
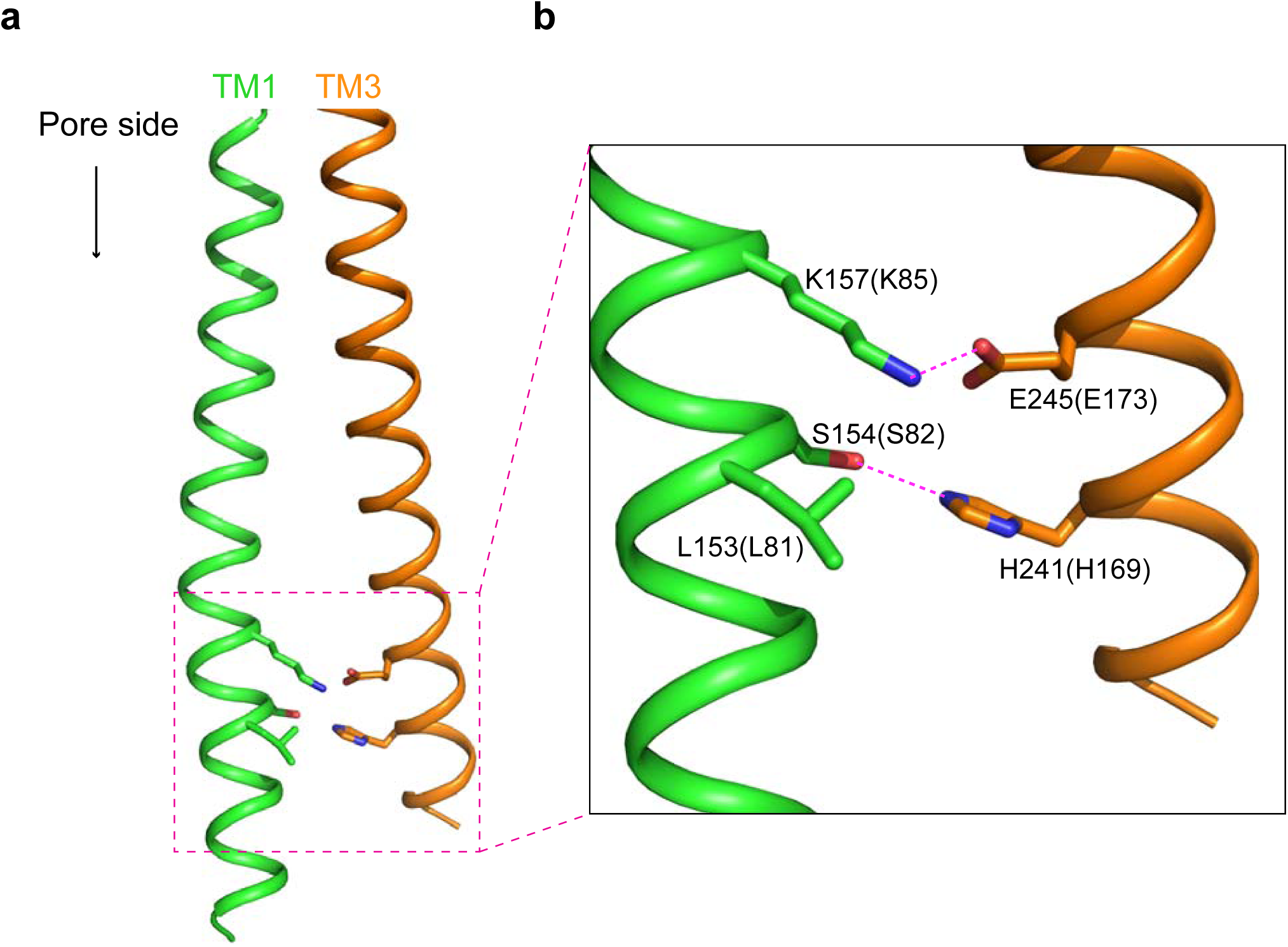
Interactions between the TM1 helix and the TM3 helix in the closed Orai structure. **a**, The TM1 helix and the TM3 helix are colored green and orange, respectively. The ion-conducting pore side is labeled. **b**, Zoom view of the specific interactions between two helices. Side chains of three residues (K157, S154 and L153) from the TM1 helix and two residues (E245 and H241) from the TM3 helix are shown. The hydrogen bonds are shown as magenta dashed line. Atoms oxygen and nitrogen are colored red and blue, respectively. Amino acids in parentheses denote human Orai1 counterparts. The atom coordinates were taken from the structure with the RCSB code 4HKR. Side chains of residues K157 and L153 were absent in original PDB file. They were manually built from the program Coot based on frequently used rotamers.

**Supplemental Figure 10.**
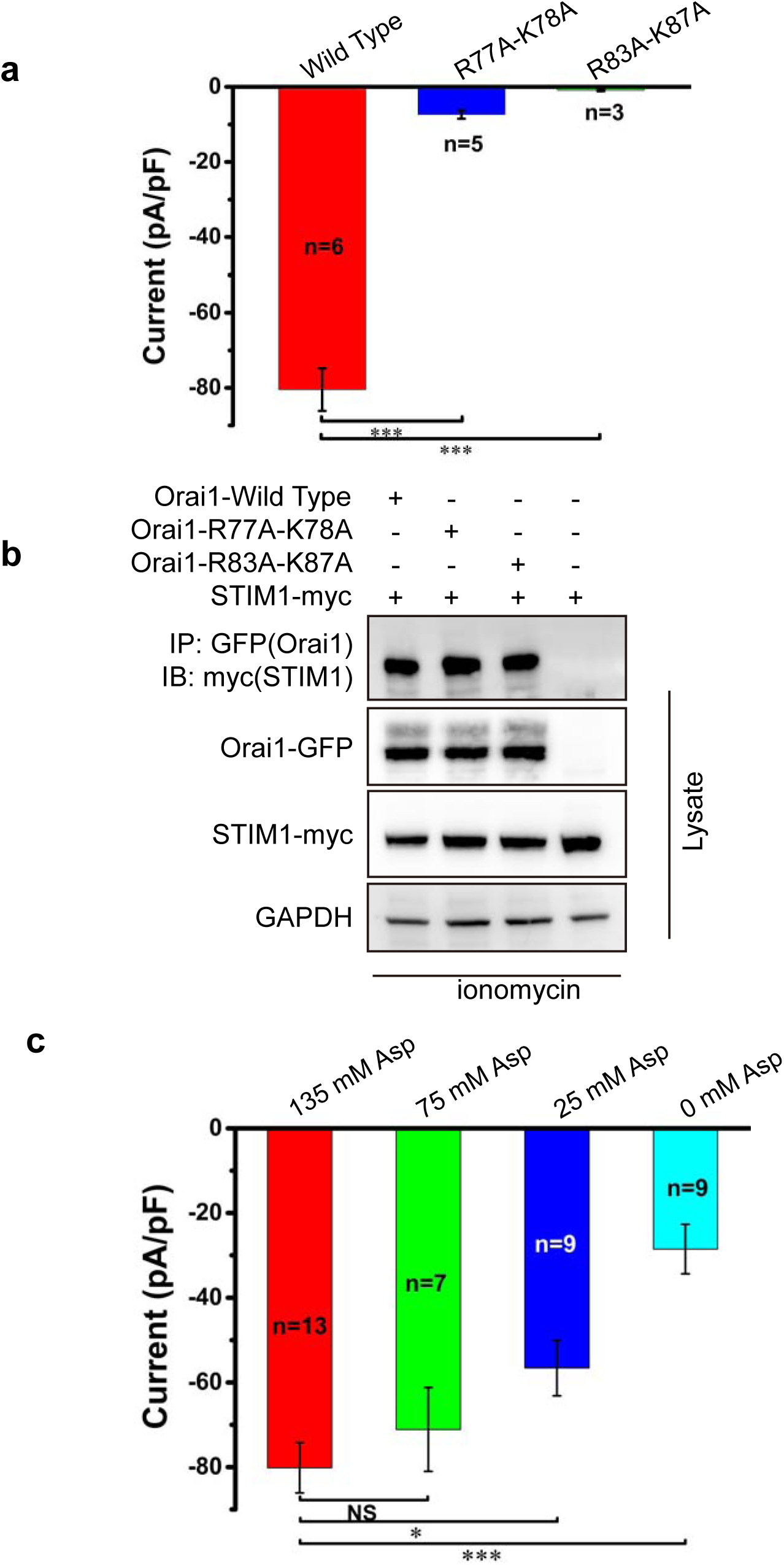
Anions near the pore on the cytosolic side are critical for Ca^2+^ permeation. **a,** Bar graphs of whole-cell Ca^2+^ currents of wild-type and mutant STIM1-activated human Orai1 channels (Orai1, Orai1-R83A-K87A and Oai1-R77A-K78A). **b,** Western blot analysis of coimmunoprecipitated human Orai1-GFP (wild type and mutants) with STIM1-myc. **c,** Bar graphs of whole-cell Ca^2+^ currents of wild-type STIM1-activated human Orai1 channels with sodium aspartate at concentrations of 0, 25 mM, 75 mM and 135 mM in the pipette solution. The number of analyzed cells is indicated. * p<0.05; *** <0.001 (unpaired Student’s t-test). Error bars denote S.E.M.

